# Generation of human cerebral organoids with a structured outer subventricular zone

**DOI:** 10.1101/2023.02.17.528906

**Authors:** Ryan Walsh, Elisa Giacomelli, Gabriele Ciceri, Chelsea Rittenhouse, Maura Galimberti, Youjun Wu, James Muller, Elena Vezzoli, Johannes Jungverdorben, Ting Zhou, Roger A Barker, Elena Cattaneo, Lorenz Studer, Arianna Baggiolini

## Abstract

Mammalian outer radial glia (oRG) emerge as cortical progenitor cells that directly support the development of an enlarged outer subventricular zone (oSVZ) and, in turn, the expansion of the neocortex. The *in vitro* generation of oRG is essential to model and investigate the underlying mechanisms of human neocortical development and expansion. By activating the STAT3 pathway using LIF, which is not produced in guided cortical organoids, we developed a cerebral organoid differentiation method from human pluripotent stem cells (hPSCs) that recapitulates the expansion of a progenitor pool into the oSVZ. The structured oSVZ is composed of progenitor cells expressing specific oRG markers such as *GFAP, LIFR, HOPX*, which closely matches human oRG *in vivo*. In this microenvironment, cortical neurons showed faster maturation with enhanced metabolic and functional activity. Incorporation of hPSC-derived brain vascular LIF-producing pericytes in cerebral organoids mimicked the effects of LIF treatment. These data indicate that the cellular complexity of the cortical microenvironment, including cell-types of the brain vasculature, favors the appearance of oRG and provides a platform to routinely study oRG in hPSC-derived brain organoids.

## Introduction

During brain development, the progenitors of the human neocortex are organized into distinct proliferative compartments. These are the ventricular zone (VZ) and the subventricular zone (SVZ), which give rise to outer neuronal layers in the cortical plate. The VZ and SVZ contain different classes of neural progenitors: apical radial glia (aRG) in the VZ, and basal, also known as outer radial glia (oRG), intermediate progenitors, and transient amplifying cells in the SVZ (Florio and Huttner, 2014). Like aRG, oRG self-renew and give rise to cortical neurons via asymmetric cell divisions (Shitamukai and Matsuzaki, 2012). Unlike aRGs, oRGs do not possess an apical process, nor do they undergo dynamic interkinetic nuclear migration within the VZ (Fietz et al., 2010; Hansen et al., 2010). In gyrencephalic species (e.g., human, macaque, ferret) the SVZ is vastly expanded and subdivided into the inner SVZ (iSVC) and outer SVZ (oSVZ). oRG primarily populate the oSVZ and oRG are largely absent in the developing cerebral cortex of lissencephalic species (e.g., mouse, rat, rabbit), linking the expansion of this compartment to neocortical expansion and folding (Lewitus et al., 2013; Reillo et al., 2011; Sun and Hevner, 2014). In current guided cerebral organoid differentiation protocols in which hPSCs are directed towards cerebral cortical identity via small molecule treatment (Pasca et al., 2022; Pollen et al., 2019; Sloan et al., 2017), oRG are less prominent and often lack the expression of the specific marker Glial Fibrillary Acid Protein (GFAP). GFAP is a comprehensive marker for glial lineage cell types including oRG (Bignami and Dahl, 1977; Johnson et al., 2016; Zhang, 2001); however, GFAP expression appears only very late in hPSC-based 3D culture (> 90 days) (Camp et al., 2015; Pollen et al., 2019; Rosebrock et al., 2022; Sloan et al., 2017; Uzquiano et al., 2022; Velasco et al., 2019). Unguided or “minimally guided” cerebral organoid differentiation protocols from hPSCs eventually lead to the spontaneous production of oRGs (Pellegrini et al., 2020), though the factors that promote the emergence of oRG and the contribution of this population to the brain microenvironment are currently poorly understood. The generation of authentic oRG is thus essential to recapitulate proper human-specific neocortex development and to study mechanisms underlying human neocortical expansion *in vitro* using cerebral organoid models.

Transcriptional profiling studies for oRG suggested that elevation of leukemia inhibitory factor receptor (LIFR)/STAT3 signaling may contribute to oRG generation or expansion in the human cortex (Pollen et al., 2015). LIF is known to promote neural stem cell (NSC) self-renewal in the human adult brain by promoting the proliferation or formation of glial progenitors in the SVZ (Bauer and Patterson, 2006); Moreover, neural progenitor cell (NPC) appearance was enhanced by LIF treatment in mouse embryoid bodies (He et al., 2006), and the addition of LIF to *in vitro* cerebral organoids elevated pSTAT3 staining and led to both a >3−fold increase in oRG and thickening of the SVZ (Watanabe et al., 2017). However, it remains unclear how transcriptionally similar LIF-induced oRG are to the fetal human counterpart, what their impact is on the organoid development, and what cells within the brain microenvironment regulate LIF expression and signaling.

Here, we developed an efficient, robust, and reproducible cerebral organoid differentiation method in which a structured oSVZ is present and abundant oRG arise upon LIF treatment. Transcriptional analysis showed that hPSC-derived oRG closely match fetal oRG. We further found that LIF promotes the derivation of molecularly distinct RG based on *GFAP* and *HOPX* expression, and that, in this context, cortical neurons are metabolically more active, mature faster, and show enhanced Ca^2+^ handling. Finally, incorporation of LIF expressing, hPSC-derived brain vascular pericytes-into *in vitro* cerebral organoids was able to partially mimic the effects observed upon LIF treatment, suggesting that brain vascular cells might partially contribute to the normal oSVZ and oRG development in humans. Taken together, this study provides a reliable *in vitro* cerebral organoid system for studying human-specific neocortex development and for investigating the mechanisms involved in human neocortical expansion.

## Results

### LIF promotes the formation of oRG-like cells in *in vitro* human cerebral organoids

To develop an organoid model that incorporates oRG more efficiently, without interfering with neuronal differentiation, we tested the impact of LIF treatment, using a cerebral cortical differentiation protocol that we developed previously (Cederquist et al., 2019). Exogenous LIF treatment was selected as candidate strategy for oRG generation given the robust expression of LIFR in oRG (Pollen et al., 2015) and lack of LIF ligand expression in cortical patterned organoids (Figure S2). Neural induction was performed using dual SMAD inhibition in combination with WNT inhibition, as described in (Rosebrock *et al*., 2022), on days 0-5, followed by dual SMAD inhibition alone form days 5-8 (Figure 1A). On day 8, two different conditions were compared: 1) standard differentiation towards neural and neuronal cell fate (control condition,(Cederquist *et al*., 2019)); and 2) treatment with human LIF from day 8 onwards to promote oRG fate (Figure 1A). To test whether LIF had the ability to promote the appearance of an oSVZ in our culture conditions, we performed immunocytochemistry at an intermediate time point within the differentiation period (day 60) and used antibodies against SOX2, TBR1 and TBR2 to visualize the rosette structures in control and LIF-treated organoids (Figure 1B). Similar to the results reported by (Watanabe *et al*., 2017), LIF treatment led to a thickening of the SVZ. We also noted the emergence of an oSVZ in cerebral organoids treated with LIF (Figure 1C), whereas VZ size and organization was comparable between conditions.

**Figure 1.**
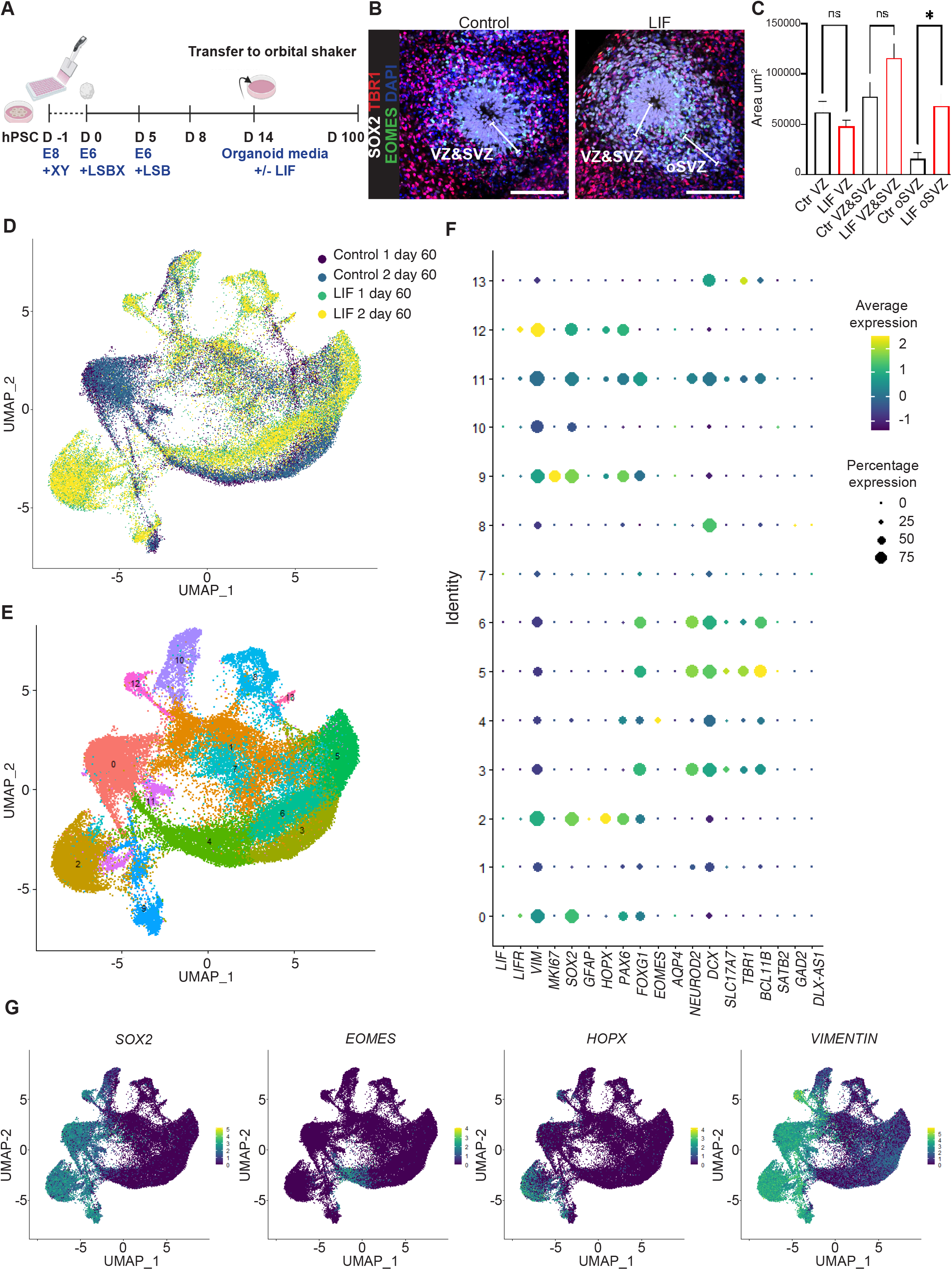
**A** The cerebral organoid differentiation protocol with or without LIF treatment. XY corresponds to XAV939 (5 μM), ROCK inhibitor (Y-27632, 10 μM); LSBX corresponds to LDN193189 (100 nM), SB431542 (10 μM) and XAV939 (5 μM); LSB corresponds to LDN193189 (100 nM) and SB431542 (10 μM); for organoid media composition see Materials & Methods. Human LIF was added at 10 ng/mL. Medium was changed every other day throughout the protocol up to day 14. From day 14, organoids were transferred to an orbital shaker in 10 cm dishes and medium was changed on a Monda-;Wednesday-Friday schedule. Half of the medium was changed throughout the protocol. **B** SOX2, TBR1 and TBR2 staining showing ventricular zone (VZ), subventricular zone (SVZ) and outer subventricular zone (oSVZ) in fixed control vs LIF- treated organoids, sectioned and stained on day 60. Nuclei are stained in blue with Dapi. Scale bars 100 μm. **C** Rosette area quantifications based on IF images from B. N=7-60 ventricles/condition across 3 independent differentiations. One-way ANOVA with Turkey’s test. Adjusted p values Ctr VZ vs LIF VZ: ns=0.966. Adjusted p values Ctr VZ&SVZ vs LIF VZ&SVZ: ns=0.240. Adjusted p value Ctr oSVZ vs LIF oSVZ: * = 0.034. Control organoids are depicted in black; LIF treated organoids are depicted in red. **D** Single-cell RNA-seq (scRNA-seq) experiments showing plots for original identity and **E** Seurat clusters in control and LIF- treated organoids on day 60. Two independent batches of 10 organoids each were processed and analyzed by scRNA-seq for both control and LIF conditions. **F** Genes selected from E to mark selected populations of interest (neuronal cells, neuronal progenitors, astrocytes, and progenitor cells including oRG) in control and LIF-treated organoid clusters from day 60. **J** Feature plots depicting distribution of the expression of *SOX2, EOMES, HOPX* and *VIMENTIN* in control and LIF-treated organoids on day 60.

Next, we investigated the transcriptional profile of control versus LIF-treated organoids using single-cell RNA-sequencing (scRNA-seq) on day 60 (Figure 1D, E). We checked the expression of transcripts related to glycolysis and endoplasmic reticulum (ER) stress, as they have been previously shown to be more predominantly expressed in oRG in organoids than in primary oRG (Bhaduri et al., 2020), but no significant difference between LIF-treated and control cortical organoids was detectable (Figure S1D). However interestingly, expression of *LIFR* was confirmed in both control and LIF-treated organoids (Figure S1A), and gene expression patterns and sample distribution revealed a molecularly distinct LIF-specific progenitor/oRG-like population characterized by the expression of typical progenitor and oRG markers such as *SOX2, HOPX* and *VIMENTIN* (Figure 1F, G; Figure S1B, C). *HOPX* was previously identified as a marker of human oRG (Pollen et al., 2015). Moreover, these cells were positive for *PAX6* but negative for *TBR2* which is consistent with an oRG identity as defined by (Matsumoto et al., 2020). Our data revealed an expansion of *HOPX*^*+*^ cells in the presence of LIF, suggesting the formation of progenitor cells with possible oRG-like identity. This *HOPX*^*+*^ population was absent on day 40 (Figure S3A-D), suggesting that oRG emerged only after day 40, but prior to day 60 under these culture conditions. Upon scRNA seq and IF analysis, we noted a LIF treatment-specific expression of *GFAP* within *SOX2*^+^ progenitor populations, including the *HOPX*^*+*^ population (Figure S1B). Of note, this *HOPX*^*+*^*GFAP*^*+*^ population also expressed the oRG markers *LIFR* and *PTPRZ1* (Figure S1D), whereas the astrocyte marker *AQP4* was not expressed (Figure 1F; FigureS3E), confirming RG rather than astrocytic identity. Interestingly, although most cells expressing *HOPX* and *GFAP* belonged to the LIF-treated condition, we noted a population of *SOX2*^*+*^*VIMENTIN*^*+*^ progenitor cells in the control population that largely lacked *HOPX* and *GFAP* expression, but similarly expressed *LIFR* (Figure S1A). scRNA-seq on day 100 revealed that a few *HOPX*^+^ cells appeared in control organoids, but these cells remained largely negative for *GFAP* (Figure S4A-D). Taken together, these findings suggest that our control condition generates a putative pre-oRG population of progenitors expressing *LIFR* that, without the inductive signal from LIF, remain functionally distinct from *HOPX*^*+*^ *GFAP*^*+*^ oRG-like cells.

### *In vitro* LIF-induced oRG-like cells are transcriptionally similar to *in vivo* oRG

In line with the scRNA-seq data, in the absence of LIF there was barely any GFAP expression by immunofluorescence (IF) staining on day 65 (Figure 2A). In contrast, in the presence of LIF, high levels of VIMENTIN and GFAP expression were detected in both the VZ and expanded oSVZ (Figure 2A). GFAP expression was still largely absent in control organoids at day100, whereas LIF-treated organoids showed abundant GFAP expression, which was consistent across multiple hPSC lines, demonstrating reproducibility of the protocol (Figure S2C-F and S4E).

**Figure 2.**
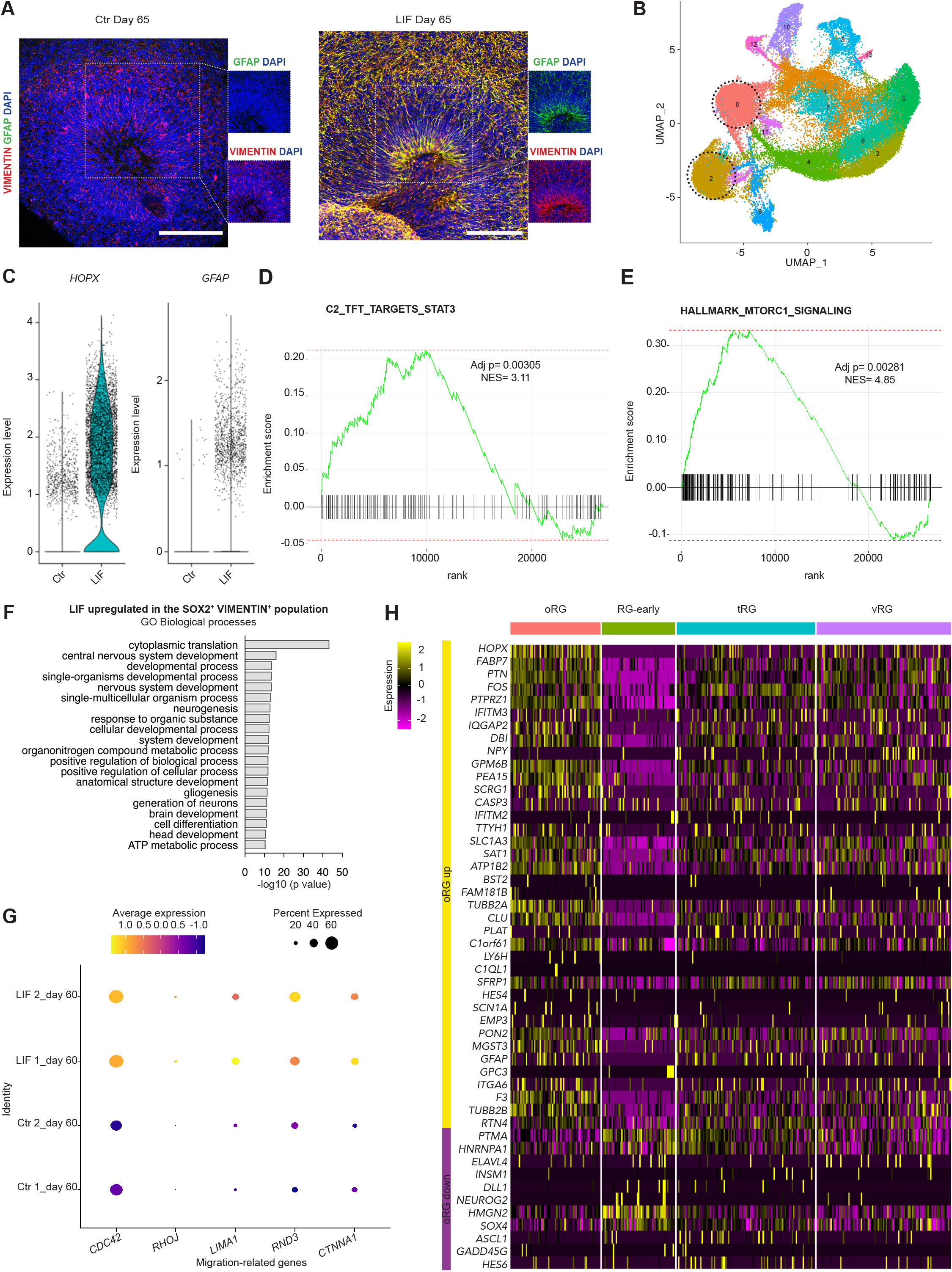
**A** GFAP and VIMENTIN staining in control (left) and LIF-treated (right) organoids that were fixed, sectioned and stained on day 65. Nuclei are stained in blue with Dapi. Scale bars:100 μm. **B** scRNA-seq experiments showing plots for Seurat clusters in control and LIF-treated organoids on day 60 from Figure 1D, E. The progenitor and oRG clusters 0 and 2 respectively are highlighted. Two independent batches of 10 organoids each were processed and analyzed by scRNA-seq for both control and LIF conditions. **C** Violin plots showing expression levels of *HOPX* and *GFAP* at day 60. **D** Enrichment analysis showing upregulation of STAT3- and **E** mTORC1 signaling in clusters 2 vs. 0 in LIF-treated organoids. **F** GO Biological Process terms enriched in clusters 2 vs. 0 in LIF-treated organoids. **G** Migration-related genes selected from F in day 60- control and LIF-treated organoids upregulated in clusters 2 vs. 0. **H** Heatmap depicting RG cell clustering distribution in LIF-treated organoids on day 60 compared to (Nowakowski *et al*., 2017). The heatmap shows the 50 most significant DE genes in *HOPX*+ vs *HOPX*-in *VIM*ENTIN+ clusters within the LIF treatment condition.

We then aimed to characterize the transcriptional profile of the *GFAP*^*+*^ *HOPX*^*+*^ cells. Within the *SOX2*^*+*^*VIMENTIN* ^*+*^ population, we compared cluster 0 (*GFAP* ^*−*^ *HOPX* ^−^) with cluster 2 (*GFAP* ^*+*^ *HOPX* ^*+*^) on day 60 (Figure 2B-F and Figure S1F). These clusters are highlighted in Figure 2B. Most of the cells expressing *HOPX* and *GFAP* belonged to LIF-treated organoids (Figure 2C). When we compared these clusters within LIF-treated cells, we found an upregulation of STAT3 and mTORC1 signaling, which is consistent with data from primary oRGs in human fetal tissue studies (Andrews et al., 2020; Pollen *et al*., 2015) (Figure 2D, E). We also observed a strong enrichment of oxidative phosphorylation system-(OXPHOS) and mitochondrial genes as well as transcripts associated with migration in cluster 2 compared to clusters 0 (Figure 2F, G). Expression of selected migration genes of interest are depicted in Figure 2G and it was higher in LIF-treated organoids compared to controls (Figure 2G). We then asked whether our oRG-like cells had a similar molecular identity to *in vivo* oRG (Figure S1E). We specifically looked at LIF-treated cells and, within the *SOX2*^*+*^ *VIMENTIN*^*+*^ population, we selected the top 50 genes based on the corrected p-value in the *HOPX* ^+^ vs. *HOPX* ^−^ cells to generate an oRG expression signature for our dataset (Figure S1F). When comparing those gene sets with *in vivo* scRNA-seq data from (Nowakowski et al., 2017), we found that our oRG signature was enriched specifically in human fetal oRG (Figure 2H), indicating that LIF-induced oRG-like cells more closely match the transcriptional profile of their *in vivo* counterparts. Of particular interest, the *in vitro HOPX* ^+^ oRG also expressed high levels of *PTPRZ1* (Figure 2H, Figure S1D), a cell-surface marker of the oRG population (Pollen *et al*., 2015), that was reported to be expressed in *HOPX* ^*+*^cells *in vivo* when compared to human cortical organoids *in vitro* (Bhaduri *et al*., 2020) further suggesting that our *GFAP*^*+*^ *HOPX*^*+*^ cells reflect authentic oRG.

### LIF treatment results in neuronal maturation in the human cerebral organoids

We observed that LIF supplementation promoted the expansion of an oRG pool without interfering with neuronal differentiation (Figure S3D, E; Figure S4D). We next focused on the neuronal cells generated in control and LIF-treated organoids and compared them transcriptionally. When looking at neuronal cell clusters (Figure 3A from Figure 1D, E), we identified cluster 5 as the most mature neuronal cluster based on marker expression (absence of *PAX6*, and highest expression of *DCX* and *TBR1*, Figure 3B-D). To sort cells along their developmental trajectory and decipher the relationships between progenitor and neuronal clusters, we performed RNA velocity analysis and defined a velocity pseudotime (Bergen et al., 2020). Pseudotime analysis revealed that neurons from LIF-treated organoids were on average farther along their pseudotime trajectory than those of control organoids (Figure 3E, F), suggesting that neurons in LIF-treated organoids were maturing faster. In line with gene expression profile, cluster 5 appeared to be the most mature neuronal cluster based on pseudotime analysis (Figure 3E). Focusing on cluster 5, Gene Ontology (GO) analysis showed enrichment in terms for mitochondrial genes and aerobic respiration in LIF-treated organoids vs. controls (Figure 3G). Expression of selected OXPHOS genes across conditions is depicted in Figure 3H. In line with these observations, we noticed a profound increase of *MEF2C* specifically in neurons from LIF-treated organoids (Figure 3I). This effect was even stronger in cluster 5 (Figure 3J). As this factor was identified as upregulated in neurons in *in vivo* transplanted organoids compared to *in vitro* controls (Bhaduri *et al*., 2020) this data suggests that LIF treatment could lead to a more “*in vivo*-like” organoid. Finally, we investigated Ca^2+^ signaling in control vs. LIF-treated organoids and found that LIF treatment promoted an increase in Ca^2+^ amplitude and calcium frequency (Figure 3K, L, Movie S1 and S2), a sign that excitability and functional maturation may be enhanced following LIF supplementation.

**Figure 3.**
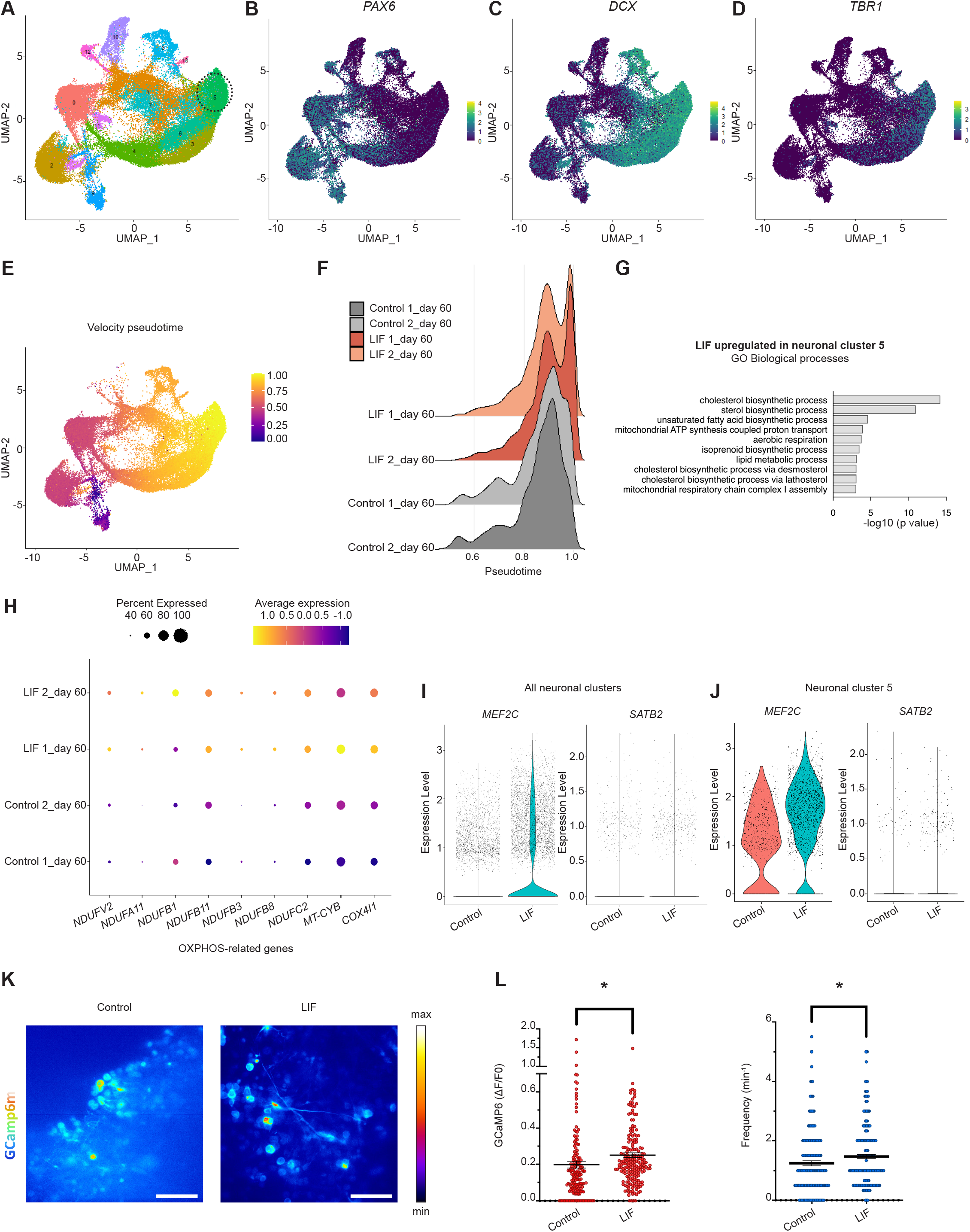
**A** scRNA-seq experiment showing plots for Seurat clusters in control and LIF-treated organoids on day 60 from Figure 1D, E. The neuronal cluster 5 is highlighted. Two independent batches of 10 organoids each were processed and analyzed by scRNA-seq for both control and LIF conditions. **B** Feature plots depicting distribution of the expression of *PAX6*, **C** *DCX* and **D** *TBR1* in control and LIF-treated organoids on day 60. **E** Velocity pseudotime plots showing all neuronal cells undergoing maturation. **F** Ridge plots showing expression levels of neuronal cells that undergo maturation in LIF-treated organoids vs. control conditions on day 60. **G** Gene Ontology (GO) Biological Process terms involved in metabolism enriched in cluster 5 and upregulated upon LIF treatment Genes selected from G to mark OXPHOS genes in control and LIF-treated organoid cells in cluster 5 on day 60. **I** Violin plots showing expression levels of *MEF2C* and *SATB2* in control and LIF-treated organoids on day 60 in all neuronal cell clusters and **J** in neuronal cell cluster 5. **K** Calcium imaging and **L** analysis showing quantification of Ca^2+^ intensity (left, Welch’s test and p value = 0.043) and frequency (right, Welch’s test and p value=0.027) measured in control and LIF-treated organoids on day 60. Scale bars: 50 μm.

### Vascular pericytes promote the emergence of oSVZ and oRG in *in vitro* human cerebral organoids, mimicking LIF treatment

To understand whether a cell type that expresses LIF could be added to cortical organoids to obviate the need for exogenous LIF, we asked how LIF ligand is produced in the brain. In other organs such as the heart, vascular pericytes are known to secrete LIF (Su et al., 2021). In the brain, vascular pericytes, that originates either from mesodermal progenitors or the neural crest (NC), play a prominent role in numerous vascular functions including regulation of cerebral blood flow, maintenance of the blood brain barrier (BBB), and control of vascular development and angiogenesis (Brown et al., 2019). NC-derived pericytes also facilitate neuroinflammatory processes, possess stem-like properties, and form part of the neurovascular unit (NVU), a group of cells that control interactions between neurons and the cerebral vasculature to meet demands of the brain (Brown *et al*., 2019). To investigate LIF expression in brain pericytes, we analysed human fetal brain sections at 11GW using pericyte markers (PDGFRb, aSMA) in combination with an anti-LIF antibody and found LIF-positivity in pericytes of the choroid plexus, a brain region that contains abundant pericytes (Figure 4A; Figure S4F). This is particularly interesting considering LIF expression in unguided brain organoids that form choroid plexus and that also show the emergence of a *GFAP*^*+*^ *HOPX*^*+*^ positive oRG-like population (Figure S2) (Pellegrini *et al*., 2020). Next, we aimed to investigate whether incorporating NC-derived pericytes in our control organoids could rescue the limited oSVZ and oRG development, thus mimicking LIF treatment. Firstly, we developed a 2-step protocol to differentiate hPSCs into NC-derived pericytes using a NC induction protocol developed previously (Baggiolini et al., 2021; Lee et al., 2010; Mica et al., 2013) followed by a pericyte differentiation method based on (Faal et al., 2019) but without the addition of serum (Figure 4B). We used the SOX10^eGFP/w^ hESC line in which enhanced green fluorescent protein (eGFP) is targeted to the genomic locus of the NC transcription factor *SOX10* (Kim et al., 2003). This allows the appearance and enrichment of NC cells to be monitored using eGFP expression. Neural crest induction was performed using a previously established protocol (Tchieu et al., 2017) based on TGFβ inhibition combined with low dose BMP and WNT activation via glycogen synthase kinase-3B (GSK3b) inhibition on days 0-2, followed by increased levels of WNT activation from days 2-10 (Figure 4B). On day 10, hPSCs-derived NC cells were sorted based on eGFP expression to derive SOX10^+^ NC cells (Figure 4B). SOX10^+^ cells were maintained and passaged using a serum-free pericyte medium for one week (Figure 4B). On day 17, cells displayed typical pericyte morphology (Figure 4C), and expressed aSMA (Figure 4D) and TAGLN by IF (Figure 4E), but not the endothelial cell marker CD31 (Figure 4F). Expression of LIF protein was detected in our NC-derived pericytes (Figure 4G). Secondly, day-17 NC-derived pericytes were aggregated into spheroid microtissues in V-bottom 96-well microplates as depicted in Figure 4H, similar to what has been described previously for microtissues containing cardiac myocytes (Giacomelli et al., 2017). Each pericyte microtissue was made up of 10,000 cells which self-aggregated within 24 hours. Pericyte microtissues were then fused to control cerebral organoids to generate cortical-pericyte (CP) assembloids as depicted in Figure 4I. Control and LIF-treated organoids were differentiated in parallel as negative and positive control, respectively. Visual assessment of rosette areas (Figure 4J) and immunocytochemistry (Figure 4K) revealed rosette expansion and oSVZ appearance in CP assembloids vs. control organoids, similarly to LIF-treated organoids (Figure 4K). Next, we compared the transcriptional profile of CP assembloids vs control and LIF-treated organoids by scRNA-seq (Figure 4L, M) and found a similar oRG cluster containing cells expressing *GFAP* and *HOPX* in CP assembloids (Figure 4N, O), Expression of pericyte markers was still maintained in a population of cells within the CP assembloid (cluster 17, Figure S4G, H). Overall, these structural and transcriptional results indicated that inclusion of hPSC-derived brain pericytes in *in vitro* cerebral organoids was able to mimic the exogenous LIF treatment and promoted oSVZ and oRG expansion.

**Figure 4.**
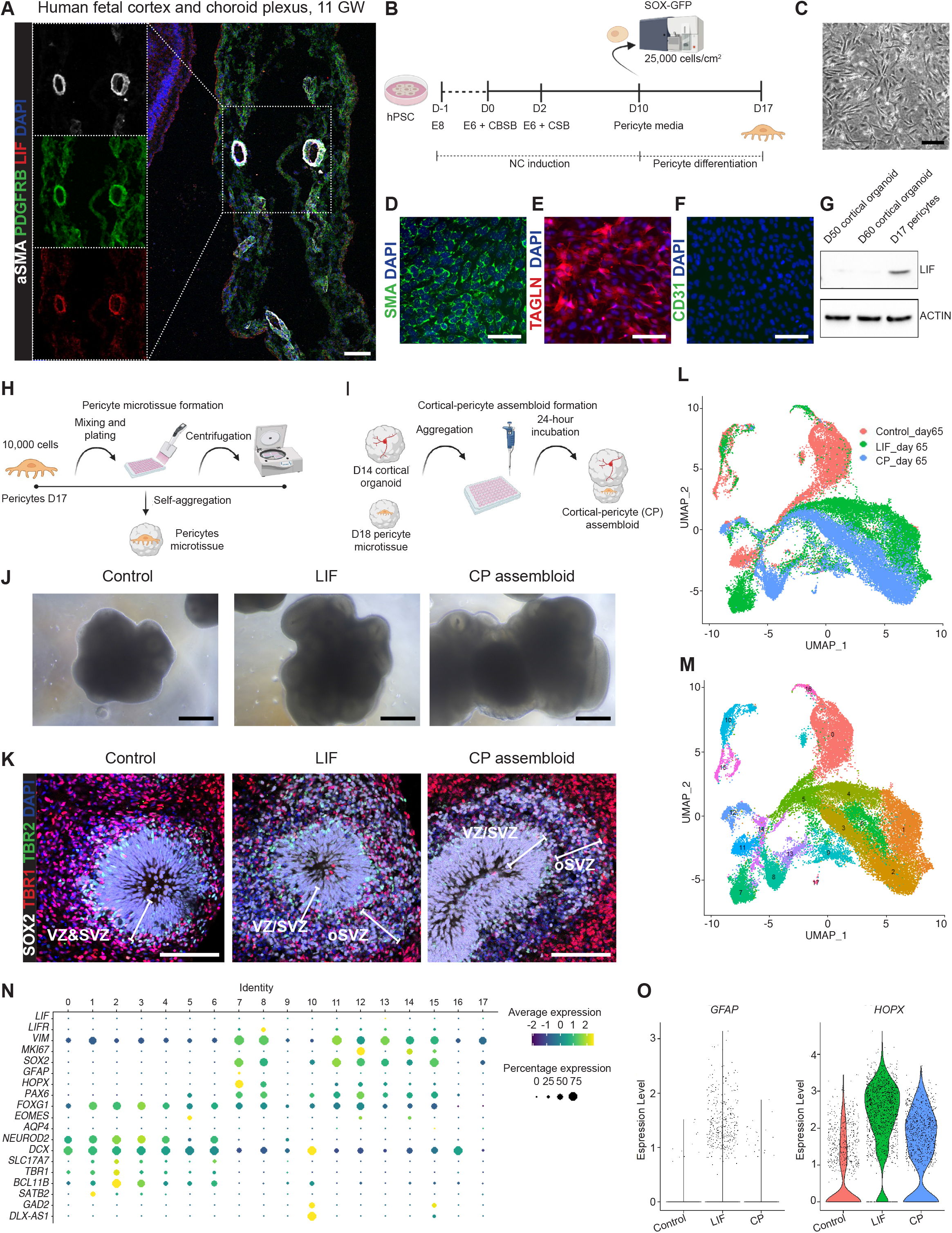
**A** LIF, PDGFRb and aSMA staining in human fetal cortex and choroid plexus from human fetal brain 11GW sections. Nuclei are stained in blue with Hoechst. Scale bars: 100 μm. **B** The pericyte differentiation protocol from H9-SOX10-GFP hPSCs-derived neural crest cells. Neural induction was performed using the dual SMAD inhibition protocol. CSBB corresponds to CHIR (600nM), SB (10μM) and BMP4 (1 ng/ml). CSB corresponds to CHIR (1.5μM) and SB (10μM). Medium was refreshed every day throughout the differentiation protocol up to day 10. On day 10, hPSCs-derived neural crest cells were sorted based on GFP to derive SOX10^+^ neural crest cells. SOX10^+^ cells were differentiated into pericyte-like cells by seeding 25,000 cells/cm^2^ single cells onto matrigel-coated plates. Cells were maintained in pericyte medium (ScienCell) without FBS for one week to generate neural crest-derived pericytes. **C** Representative bright-field images of the morphological appearance of neural crest-derived pericytes from H9-SOX10−GFP hPSCs on day 17. Scale bars: 100 μm. **D** immunofluorescence images of TAGL, **E** SMA and **F** CD31 in neural crest-derived pericytes from H9-SOX10-GFP hPSCs on day 17. Pericytes stain positively for TAGLN and SMA, but negatively for the endothelial marker CD31. Scale bars: 100 μm. **G** Western blot data showing LIF expression in day 17-pericytes but not in control organoids (on day 50 and 60). Pericytes were differentiated from H9-SOX10-GFP hPSCs whereas organoids came from H9 hPSCs. **H** Schematic showing the protocol to form pericyte microtissues and **I** cortical-pericyte (CP) assembloids. On day 17, hPSC-pericytes were detached and resuspended in pericyte medium. Pericytes were diluted to 10,000 cells per 100 μl of medium. Cell suspensions were seeded on V-bottom 96 well microplates and incubated at 37°C, 5% CO2 for 24 hours prior cortical-pericyte assembloid formation. Day 14-cortical organoids (without LIF treatment) and day 18-pericyte microtissues were derived separately and fused by placing them in close proximity in V-bottom 96 well microplates. Plates were centrifuged for 1 min at 1100 rpm and placed in the incubator for 24 hours. After 24-hour incubation, assembloids were moved to an orbital shaker on 10 cm dishes. Half of the medium was changed on a Monday-Wednesday- Friday schedule. **J** Representative bright-field images of the morphological appearance of control organoids, LIF-treated organoids and CP assembloids on day 60 **K** SOX2, TBR1 and TBR2 staining showing rosette areas. Scale bars: 100 μm. **L** scRNA-seq experiments showing plots for original identity and **M** Seurat clusters in control organoids, LIF-treated organoids, and CP assembloids on day 65. **N** Genes selected from N to mark selected populations of interest (dividing cells, neuronal cell types, and neuronal progenitors, astrocytes, postmitotic neurons and radial glia) in control, LIF-treated organoid and CP assembloid clusters on day 65. **O** Violin plots showing expression levels of *GFAP* and *HOPX* in control vs. LIF-treated organoids vs. CP assembloids on day 65.

Our data indicate that inclusion of brain vascular cells and potentially other non-neural derived lineages in cerebral organoid systems will be important to better recapitulate the physiological development of human cortical lineages.

## Discussion

In this study, we established a hPSC-derived platform for the induction and expansion of *GFAP*^*+*^*HOPX*^*+*^ oRGs in human cerebral organoids that transcriptionally match human fetal oRG. This was achieved by either supplementation of cortical organoids with human recombinant LIF or via incorporation of hPSC-NC-derived pericytes. Interestingly, several guided cerebral organoid differentiation protocols show a lack or a very delayed appearance of *GFAP*^*+*^ *HOPX*^*+*^ oRGs (Figure S2A, B; (Camp *et al*., 2015; Pollen *et al*., 2019; Sloan *et al*., 2017). In contrast, some unguided differentiation protocols, commonly associated with the formation of choroid plexus structures, a region that produces LIF both *in vitro* (Pellegrini *et al*., 2020) and *in vivo* (Figure S4F), seem to contain populations of *GFAP*^*+*^ *HOPX*^*+*^ oRG (Figure S2A, B).

Two distinct types of oRG were previously identified based on *HOPX* expression in the gyrencephalic cerebral cortex of ferrets; these were *HOPX*^*+*^ and *HOPX*^*−*^ oRG. Compared to *HOPX*^*−*^ oRGs, *HOPX*^+^ oRGs had higher self-renewal activity and preferentially produced upper-layer neurons in prospective gyral regions, resulting in the formation of cortical folds in gyrencephalic brains (Matsumoto *et al*., 2020). Within the *SOX2*^*+*^ *VIMENTIN*^*+*^ progenitor cells, we also identified *HOPX*^−^ and *HOPX*^*+*^ cells in LIF-treated organoids. In the future, it will be interesting to isolate these two cell populations directly from LIF- treated organoids and compare their self-renewal properties and differentiation capabilities.

The fact that our oRG express *GFAP* and *HOPX* simultaneously, in combination with the transcriptional similarity to *in vivo* oRG from (Nowakowski *et al*., 2017), strongly indicate that we obtained and expanded bona-fide oRGs. Our work highlights the importance of the microenvironment in the differentiation of NSCs into oRG. The population of *SOX2*^*+*^ *VIMENTIN*^*+*^ cells in control organoids also express *LIFR*, though they fail to express oRG specific markers. Our results indicates that the emergence of oRG requires extrinsic signals and highlight LIF as a critical factor in this process. Indeed, virtually no cells are producing LIF in current guided differentiations (Figure S2A, B, Figures S3E) and the lack of LIF prevents a sustained activation of the STAT3 pathway, which seems to be required for efficient oRG production. Interestingly, the addition of pericytes, a cell type that contributes to the formation of the blood brain barrier and produces LIF (Figure 4A), partially rescues the lack of *HOPX*^*+*^*GFAP*^*+*^ oRGs in our protocol (Figure 4), underlining the potential implication of the brain vasculature not only in sustaining brain-specific cells, but also in regulating their differentiation and cellular architecture.

In addition to vascular pericytes, other cell types express LIF in the human brain. For instance, microglia can also produce LIF and promote the differentiation of neural progenitor cells into astrocytes (Nakanishi et al., 2007). Similarly, cells in the choroid plexus are also a major source of LIF (Gregg and Weiss, 2005); and Figures S4F). Our CP assembloids offer the proof-of-principle that by increasing the cellular complexity of the organoid microenvironment and adding a cell type secreting LIF, it is possible to activate the STAT3 pathway and expand the oRG population. In line with this finding, a previously published protocol of unguided differentiation (Pellegrini *et al*., 2020) generated a subset of choroidal plexus cells expressing LIF (Figures S2B), which also contained a *GFAP*^*+*^*HOPX*^*+*^ population (Figure S2A).

The neural population in LIF-treated organoids showed an enhanced OXPHOS profile and Ca^2+^ activity. We also observed a trend towards an increased production of SATB2 neurons in the LIF-treated conditions, but further studies will be needed to confirm whether LIF treatment consistently promotes upper layer neuron formation (Figure 3). The reason why neurons in LIF-treated organoids express more mature features based on transcriptional and functional readouts (Ciceri et al., 2022) is unclear and it will be an exciting new topic of research. Similarly, we observed a trend for increased generation of cortical derived interneurons (Figure 3), but this population was eventually found also in control organoids (Figure 4), and additional work will be required to prove the specific impact of LIF on the differentiation of cortical derived interneurons and whether those cells represent oRG-derived cell types.

It is conceivable that the presence of oRG and the formation of a structured oSVZ promotes the health of the brain microenvironment, and this translates to enhanced neuronal activity. Another interesting hypothesis is that the derivatives of oRG might mature faster than those derived from aRG and future studies will be needed to shed light on these possibilities. Interestingly, OXPHOS related genes showed increased expression in LIF-treated oRG, and forced enhancement of OXPHOS has been linked to accelerated neuronal maturation (Iwata et al., 2021).

This study describes a cortical organoid system that incorporates an expanded and more consistent germinal zone (oSVZ) and progenitor population (oRG), two features characteristic for primate cortical development. Therefore, our findings may open the possibility to use this system for future mechanistic studies comparing neocortical development and expansion across different species. Finally, we demonstrate that the presence of vascular pericytes is important for recapitulating important aspects of neocortical development such as the production of oSVZ and oRG. This finding points to the importance of developing more complex organoid models that incorporate vascular and other non-neural cells to study their contribution to neocortex development as well as to cortical neuronal maturation.

## Acknowledgements

We thank all the Studer Lab members and Raffaele Luongo from the Baggiolini lab for insightful comments and feedback on this manuscript. We would also like to thank the Flow Cytometry core, the Molecular Cytology core, the Integrated Genomic Operation at MSKCC for outstanding technical support. We acknowledge the use of the Integrated Genomics Operation Core, funded by the NCI Cancer Center Support Grant (CCSG, P30 CA08748), Cycle for Survival, and the Marie-Josée and Henry R. Kravis Center for Molecular Oncology. This work was supported in part through the NIH grants R01AG054720, R21NS116545 (L.S.), the grant from the Department of Defense (DOD) AL200169 - W81XWH2110140 (L.S.) and the support from the Tri institutional stem cell initiative. A.B. was supported by the Alan and Sandra Gerry Metastasis and Tumor Ecosystems Center (GMTEC) Scholars Fellowship Program and by the Foundation for the Institute of Oncology Research. G.C. was supported by the European Molecular Biology Organization (EMBO) long-term fellowship (ALTF 311-2015) and the New York State Stem Cell Science (NYSTEM) postdoctoral training award (C32559GG), E.G. by the Rubicon (2020/30766/ZONMW) and R.W. by the NIH F32 (5F32MH116590-03). Processing of the human fetal brain specimen was funded by the European Union’s Horizon 2020 Research and Innovation Programme under grant agreement No. 874758 - Consortium Nsc-Reconstruct: Novel Strategies for Cell based Neural Reconstruction (2020-23) awarded to E.C. and R.B. R.B. was supported by the NIHR Cambridge Biomedical Research Centre (NIHR203312). The views expressed are those of the author(s) and not necessarily those of the NIHR or the Department of Health and Social Care. This research was funded in part by the Wellcome Trust 203151/Z/16/Z. For the purpose of open access, the author has applied a Creative Commons Attribution (CC BY) licence to any Author Accepted Manuscript version arising from this submission. Ethics number is 096/085.

## Author Contribution

L.S., A.B., R.W., E.G., G.C. and C.R. conceived and designed the experiments, performed data analysis and interpretation, and wrote manuscript.

E.G., G.C. and C.R. performed the hPSC differentiations into cortical brain organoids, neural crest and pericytes, and their analyses.

R.W. performed the sc-RNAseq analyses.

M.G., E.V., E.C., and R.B. provided and analyzed the human fetal cortex samples.

Y.W. and T.Z. engineered the SOX10-GFP hPSC line.

J.M., G.C., performed the live imaging of intact organoids at the light sheet microscope. J.J., performed some of the organoid characterization.

All authors provided feedback in editing the manuscript.

## GEO accession number

data submitted and accession number pending.

## Materials & Methods

### Experimental design

To ensure the reproducibility of the differentiation protocol provided, several members of the Studer lab independently performed aspects of the cerebral organoid differentiation method (with and without human LIF supplementation), and the protocol is consistently used in the laboratory across multiple hPSC lines. No specific methods were used for randomization and investigators were not blinded to the methods of differentiation. No statistical methods were utilized to determine sample size.

### hPSC lines culture

Human pluripotent stem cells, (H9; H9-S0×10-GFP; MEL1; MEL1 GSK3 KO; MEL1 CHD8 KO) were maintained in Essential 8 medium in feeder-free conditions on vitronectin (VTN-N) substrate. hPSCs were passaged as clumps with an EDTA dissociation solution (0.5 mM EDTA/PBS). Cells were maintained at 37°C and 5% CO2 and were routinely tested for mycoplasma and periodically assessed for genomic integrity by karyotyping.

### Generation of H9-SOX10-GFP reporter line

H9 SOX10-GFP reporter lines were generated using CRISPR/Cas9 based HDR (Zhong et al., 2020). Briefly,, a sgRNA was designed to target a sequence close to the stop codon of *SOX10* gene, and the target was cloned into the pX330-U6-Chimeric_BB-CBh-hSpCas9 vector (Addgene plasmid #42230) to make the gene targeting constructs. A donor plasmid containing a 450 bp left homology arm, followed by a P2A-H2B-GFP cassette, a floxed puromycin selection cassette and a 450 bp right homology arm was used as the donor template for knock-in. The sgRNA and the donor plasmid were electroporated into H9 cells using a Lonza 4D-Nucleofector instrument with *Solution* “Primary Cell P3”, and *Pulse Code* “CB-150”. 0.5 ug/ml Puromycin was added to the 3 days post-electroporation for 4 days. Single-cell clones were generated, PCR and sanger-sequencing were used to correctly identify knock-in clones.

### Cerebral organoid differentiation protocol and LIF treatment

On day -1, hPSCs were dissociated with EDTA for 10 minutes at 37°C and allowed to aggregate into spheroids of 10,000 cells each in V-bottom 96 well microplates (S-Bio) in E8 medium with ROCK inhibitor (Y-27632, 10 μM) and WNT inhibitor (XAV939, 5 μM, Tocris 3748). The next day (day 0), medium was changed to E6 supplemented with 3 inhibitors (LDN193189, 100 nM; SB431542, 10 μM; XAV939, 5 μM) with media change every other day. On day 5, medium was switched to E6 supplemented with LDN193189, 100 nM and SB431542, 10 μM, with media change every other day. On day 8, the medium was changed to an N2/B27 based organoid medium as previously described (Cederquist *et al*., 2019) +/- Human LIF (10 ng/mL) with media change every other day. On day 14, organoids were moved to an orbital shaker on 10 cm dishes. Half of the medium (+/- LIF treatment) was changed on a Monday-Wednesday-Friday schedule.

### Differentiation of hPSCs into neural crest-derived pericytes

Neural induction using the dual SMAD inhibition protocol was performed as previously described (Baggiolini *et al*., 2021; Lee *et al*., 2010; Mica *et al*., 2013). Prior to starting the differentiation, on day -1, H9-SOX10-GFP hPSCs were plated as a high-density monolayer (150,000 cells per cm^2^) on Matrigel in E8 medium with 10μM ROCKi. On day 0, the medium was changed to E6 supplemented with 1 ng/ml BMP4 + 10μM SB + 600nM CHIR. On day 2, the medium was changed to E6 supplemented with 10μM SB + 1.5μM CHIR. Medium was refreshed every day throughout the differentiation protocol up to day 10. On day 10, hPSCs-derived neural crest cells were sorted using a BD-FACS Aria6 cell sorter at the Flow Cytometry Core Facility of MSKCC to derive GFP^+^ neural crest cells (SOX10^+^ cells). 4, 6-diamidino-2-phenylindole (DAPI) was used to exclude dead cells. SOX10^+^ cells were differentiated into pericyte-like cells as described previously (Faal *et al*., 2019) by seeding 25,000 cells/cm^2^ single cells onto matrigel-coated plates. Cells were maintained in pericyte medium (1201, ScienCell) without FBS for one week to generate neural crest-derived pericytes. During that week, medium was refreshed every other day and cells were passaged once using Accutase and replated at a concentration of 25,000 cells/cm^2^ in pericyte medium without FBS.

### Pericyte microtissue formation

Pericyte microtissues were generated similar to that done previously for MT-CM microtissues (Giacomelli *et al*., 2017). On day 17, hPSC-pericytes were detached using Accutase for 5 min at 37 °C, 5% CO2, centrifuged for 3 min at 1100 rpm and resuspended in pericyte medium without FBS, but with ROCK inhibitor (Y-27632, 10 μM). Pericytes were diluted to 10,000 cells per 100 μl of medium. Cell suspensions were seeded on V-bottom 96 well microplates (S-Bio) and centrifuged for 10 min at 1100 rpm. Pericyte microtissues were incubated at 37°C, 5% CO2 for 24 hours (day 18-pericyte microtissues) prior to cortical-pericyte assembloid formation.

### Cortical-pericyte assembloid formation

To generate cortical-pericyte assembloids, day 14-cortical organoids (without LIF treatment) and day 18-pericyte microtissues were fused by placing them in close proximity in V-bottom 96 well microplates. Plates were centrifuged for 1 min at 1100 rpm and placed in the incubator for 24 hours at 37°C, 5% CO2. After 24 hours, tissues were fused and assembloids were moved to an orbital shaker on 10 cm dishes. Half of the medium (organoid medium without LIF treatment) was changed on a Monday-Wednesday-Friday schedule.

### Organoid and assembloid dissociation

10 organoids/assembloids for each condition and for each time point were dissociated for single-cell RNA-sequencing using a papain dissociation method following manufacturerin structions http://www.worthington-biochem.com/PDS/.“Sample” refers to 10 organoids/assembloids that were differentiated in the same dish on the same day. Briefly, samples were slightly minced (but not manually chopped) into smaller pieces by pipetting up and down ∼10 times with a P1000 in 1 ml of papain solution/sample in 1.5 Eppendorf tubes. Eppendorf tubes containing the tissues were incubated at 37°C with constant agitation on a rocker platform in the incubator for 1 hour and 15 minutes. After incubation, mixtures were gently pipetted up and down with a P1000 for ∼ 10 times, transferred to 15 ml tubes and centrifuged at 300g for 5 minutes at room temperature. Supernatants were discarded and cell pellets were immediately resuspended in DNase dilute albumin-inhibitor solution. 1 ml of albumin-inhibitor solution was added to each centrifuged tube (sample). Tubes were centrifuged at 70g for 6 minutes at room temperature. For single-cell RNA-sequencing, cells were resuspended in FACS buffer solution (4ml of FACS buffer + 20 μl of RiboLock RNase inhibitor) at a final concentration of 200,000 of cells / 200 μl of FACS buffer solution for each sample.

### Western blot

hPSC-derived pericytes and organoids without LIF treatment were lysed with RIPA buffer + 1:1000 Halt™ Protease and Phosphatase Inhibitor Cocktail (Thermo Fisher Scientific) and then sonicated for 3×30sec at 4°C. Supernatant was collected upon 15min of centrifugation at >15000 rpm at 4°C and quantified by Precision Red Advanced Protein Assay (Cytoskeleton). Equal amounts of protein were boiled in NuPAGE LDS sample buffer (Invitrogen) at 95°C for 5 min and separated using NuPAGE 4%–12% Bis-Tris Protein Gel (Invitrogen) in NuPAGE MES SDS Running Buffer (Invitrogen). Proteins were electrophoretically transferred to a nitrocellulose membrane (Thermo Fisher Scientific) with NuPAGE Transfer Buffer (Invitrogen). Blots were blocked for 60 min at RT in TBS-T + 5% nonfat milk (Cell Signaling) and incubated with the respective primary antibody at 4°C. The following primary antibodies were used: mouse anti bactin-HRP (abcam, ab49900, 1:10000), rat anti LIF (abcam, ab138002, 1:500). Primary antibodies were detected using the secondary anti-rat IgG HRp-linked (Cell Signaling, 7077S, 1:1000) together with the SuperSignal™ West Femto Chemiluminescent Substrate (Thermo Fischer Scientific).

### Histology and immunocytochemistry

Organoids and assembloids were fixed in 4% PFA overnight at 4°C and washed three times with PBS the next day. After fixation, tissue was cryoprotected in 30% sucrose/PBS and sectioned at 30μm on a cryostat (Leica 3050 S). Sections were blocked for 30 min in 10% FBS, 1% BSA, 0.3% triton in PBS, and incubated as floating sections or, alternatively, as slide-mounted sections, in primary antibodies overnight. The next day, sections were washed with PBS and incubated as floating sections or slide-mounted sections in secondary antibodies for 3 h at room temperature. DAPI stain was used to identify cell nuclei. Images were captured using a LeicaSP8WLL confocal laser-scanning microscope, or a standard inverted epifluorescence microscope (Zeiss Axio Observer). Antibodies used: rat anti SOX2 (Thermo Fisher, 14-9811-82, 1:200), mouse anti EOMES (Thermo Fisher, 14-4877-82, 1:500), rabbit anti TBR1 (Abcam, ab183032, 1:500), rat anti LIF (abcam, ab138002, 1:200), rabbit anti PDGRb (Invitrogen, MA5-15143, 1:100), mouse anti SMA (Sigma, A2547, 1:500)

*In vitro*, day 17-pericytes were fixed in 4% PFA for 10 min at room temperature, permeabilized with 0.3% Triton for 5 min, washed with PBS for 5 min and blocked with 10% donkey serum in PBS for 1 hour. Primary antibodies were diluted in blocking solution and incubated with the samples overnight at 4 °C. The next day, samples were washed 3 times with PBS and secondary antibodies were diluted in blocking solution and incubated with the samples at room temperature for 1 hour. Samples were washed 3 times and DAPI stain was used to identify cell nuclei. Microscopy was performed using a standard inverted epifluorescence microscope (Zeiss Axio Observer). Images were acquired using Zen Pro (Zeiss) Software. Antibodies used: Anti-Human CD31 Clone JC70A (Dako, M0823, 1:500); Monoclonal Anti-Actin, a-Smooth Muscle (Sigma Aldrich, A2547, 1:500); Anti-TAGLN/Transgelin antibody (Abcam, ab14106, 1:1000); donkey anti-mouse Alexa Fluor 488 (Thermo Fisher, A-21202, 1:1000), donkey anti-rabbit Alexa Fluor 555 (Thermo Fisher, A-31572, 1:1000)

### Human fetal brain sections and immunohistochemistry

Brains were dissected out, fixed (24 h, 4°C) in 4% paraformaldehyde (PFA) and incubated in sucrose 30% for 24 h at 4°C. Brains were then frozen in optimal cutting temperature (OCT) medium on dry ice. Frozen sections (15 μm) were obtained using a cryostat (Leica, Germany) and stored at - 80 °C. Sections were thawed just prior to staining and fixed with 4% PFA for 10 min followed by rinsing in PBS. Sections were then dehydrated in 5 min dehydration steps in 70% and 100% ethanol, respectively, washed with PBS, permeabilized with 0.5% Triton X-100 (Euroclone) in PBS for 10 min, washed in PBS and retrieved with Sodium Citrate 10mM at 90°C for 30min. After antigen retrieval the sections were washed with PBS and blocked with 10% NGS (Vector) and 0.2% Triton X-100 in PBS at room temperature for 1h. Primary antibodies were diluted in solution containing 3% NGS and 0.1% Triton X-100 in PBS at 4 °C overnight. The following day sections were washed three times in PBS at room temperature. Secondary antibodies conjugated to Alexa fluorophores 488, 568 or 647 (Molecular Probe, Life Technologies) were used 1:500 in solution containing 3% NGS and 0.1% Triton X-100 in PBS at room temperature for 1 h mixed with Hoechst 33258 (5 μg/ml; Thermo Fisher Scientific) to visualize nuclei. The sections were then washed once in PBS with 0,1% Triton X-100 and twice in PBS and finally mounted with Dako Glycergel (Aqueous Mounting Medium, Agilent) at room temperature overnight. The following day the sections were dry enough to be visualize under the microscope and then stored at 4°C. Images were acquired with a widefield and a confocal microscope (Leica SP5).

### Rosette quantification

Rosettes were quantified in FIJI by Image J. The ROI of the VZ was drawn around the region marked by SOX2+ cells. The ROI of the VZ/SVZ was drawn around the regions encompassing both SOX2+ cells (VZ) and the region marked by a mixture of SOX2+ and TBR2+ neural stem cells (SVZ). In order to determine the area of the oSVZ, marked by the expanding radial glial population, we subtracted the VZ area from the VZ/SVZ total area. The differential area between the two ROI’s is representative of the oSVZ. All quantifications were acquired in pixels and converted to um2 by normalizing the values to the image scale bar in um.

### Single-cell RNA-sequencing data processing and analysis

Single cell suspensions were diluted to 1,000 cells/μl in 1X PBS with 0.04% BSA and 0.2U/μl Ribolock RNAse inhibitor (Thermo #EO0382) for sequencing. Single-cell RNA-sequencing (scRNA-seq) was performed for a target recovery of 10,000 cells/sample using 10X Genomics Chromium Single Cell 3’ Kit, Version 3 according to the manufacturer’s protocol. Libraries were sequenced on an Illumina NovaSeq. The CellRanger pipeline (Version 6.1.2) was used to demultiplex and align reads to the GRCh38 reference genome to generate a cell-by-gene count matrix. Data analysis was performed with R v4.1 using Seurat v4.2.0 (Hao & Hao et al. 2021). Cells expressing between 200 and 5,000 genes and less than 10% counts in mitochondrial genes were kept for analysis. Gene counts were normalized by total counts per cell and ScaleData was used to regress out cell cycle gene expression variance as determined by the CellCycleScoring function. PCA was performed on scaled data for the top 2000 highly variable genes and a JackStraw significance test and ElbowPlot were used to determine the number of PCs for use in downstream analysis. A Uniform Manifold Approximation and Projection (UMAP) on the top 35 PCs was used for dimensional reduction and data visualization. FindNeighbors on the top 35 PCs and FindClusters with a resolution of 2.0 were used to identify clusters. Differential expression analysis was performed by identifying either oRG-like clusters or neuronal clusters based on marker expression and performing FindMarkers within these groups between LIF-treated and control cells using a Wilcoxon Rank Sum test. For trajectory and pseudotime analyses, Velocity v0.17.17 (La Manno et al., 2018) was used to define spliced vs unspliced reads, and loompy.combine (Loompy v2.0.17; http://loompy.org) was used to merge samples into a single loom file. Downstream analysis was carried out in Python v3.9.12 with Scanpy v1.9.1 (Wolf et al., 2018) and scVelo v0.2.4 (Bergen *et al*., 2020) scvelo.pp.filter_and_normalize(min_shared_counts =20, n_top_genes =5000, flavor = “seurat”) was used for normalization and log transformation and moments and nearest neighbors were then calculated with scvelo.pp.moments. Velocities were estimated using the stochastic model with scvelo.tl.veloctiy(mode = “stochastic”) and pseudotimes obtained with scv.tl.velocity_pseudotime.

### Calcium imaging and analysis

hPSC-derived cortical brain organoids were infected with lentiviruses encoding GCamp6m at day 50 of differentiation and cultured in BrainPhys™ Imaging Optimized Medium (Stem Cell Technologies) for a week before the imaging. On the day of the imaging, organoids were equilibrated in imaging buffer for 30 min (25 mM HEPES, 140 mM NaCl, 8 mM KCl, 1 mM MgCl2, 10 mM glucose, 4 mM CaCl2, 10 μM glycine, 0.1% BSA pH 7.4, pre-warmed to 37 °C) and transferred into imaging cuvettes. GCamp6m fluorescence on intact organoids was recorded by light-sheet microscopy on TruLive3D Imager (Bruker) under environmental control (37°C; 95% O2 – 5% CO2). Multiple fields of view from 3-4 organoids per condition were imaged for ∼2 min at a frame rate of 10 frames/second (1200 frames/time lapse) and at 31.3x effective magnification.

Analysis was performed as previously described (Ciceri *et al*., 2022; Sun and Südhof, 2021). Briefly, the live-imaging image stack was converted to TIFF format and loaded into optimized scripts in MATLAB. Neurons were identified based on GCaMP6m expression and regions of Interest (ROI) were placed on the neuron somas to calculate the raw GCaMP6m intensity of each neuron over time. The signal intensity of each trace was normalized to the baseline (ΔF/F0) for spike detection. The amplitude of Ca^2+^ spikes was calculated from the normalized GCaMp6m intensity for all the detected spikes in each trace (mean ΔF/F0 of detected spikes for each neuron). The frequency of Ca^2+^ spikes was calculated as the number of detected spikes in each trace per minute of recording.

### Statistics and reproducibility

All data presented in this study are representative of at least three independent experiments. Western blots were repeated at least twice. No statistical methods were used to predetermine sample sizes, but our sample sizes are similar to those reported in previous publications. Data distribution was assumed to be normal, but this was not formally tested. Data are presented as the mean ± sem. All statistical analyses were performed using GraphPad Prism 9: one-way ANOVA with Turkey’s tests to compare multiple groups and t-test to compare two groups. Statistical differences were considered significant with P < 0.05 as indicated in figure legends. All reported measurements are from distinct samples. For scRNA-seq experiments, two independent batches of 10 organoids each were processed and analyzed by scRNA-seq for both control and LIF conditions on day 40, 60 and 100, whereas one batch of 10 organoids were processed and analyzed for control, LIF condition and CP assembloids on day 65.

---

**Figure S1.**
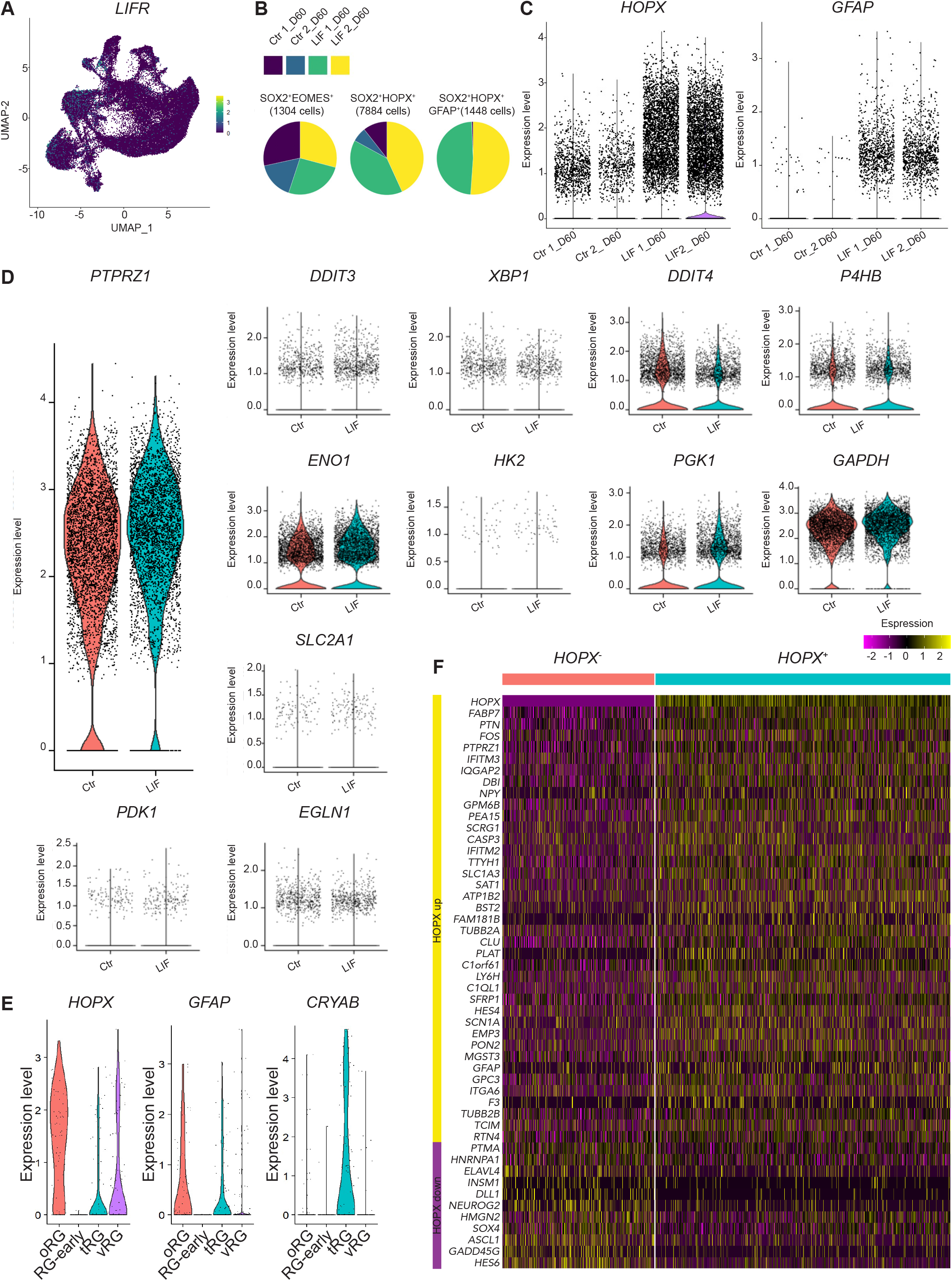
**A** Feature plot depicting distribution of the expression of LIF receptor (*LIFR*) in control and LIF-treated organoids on day 60. **B** Pie charts showing distribution of *SOX2*^*+*^*EOMES*^*+*^, *SOX2*^*+*^*HOPX*^*+*^ and *SOX2*^*+*^*HOPX*^*+*^*GFAP*^*+*^ cells across samples. **C** Violin plots showing expression levels of *HOPX* and *GFAP* in control and LIF-treated organoids on day 60 **D** Violin plot showing expression levels of *PTPRZ1* as well as glycolysis and endoplasmic reticulum (ER) stress-related transcripts in control and LIF-treated organoids on day 60 **E** Violin plots showing expression levels of *HOPX, GFAP* and *CRYAB* from (Nowakowski *et al*., 2017) **F** Hierarchical clustering heatmap analysis of control and LIF-treated organoids on day 60 showing HOPX^+^ and HOPX^-^ clusters of gene expression within the *SOX2*^*+*^ *VIMENTIN*^*+*^ population. The top 50 DE genes by p value.

**Figure S2.**
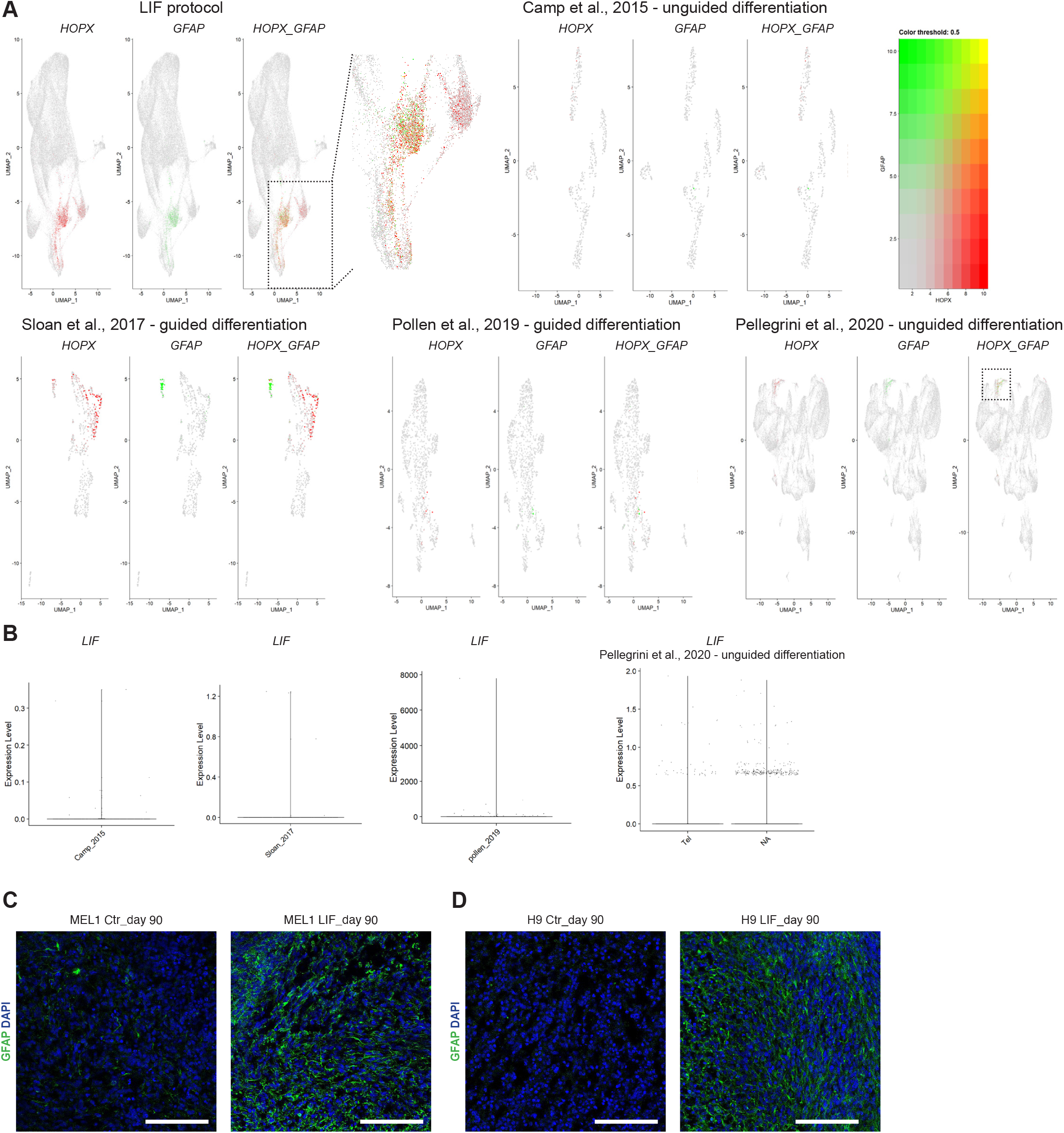
**A** Feature plots depicting distribution of *HOXP, GFAP*, and co-expression of *HOXP* and *GFAP* from our study compared to other cerebral organoid studies **B** Expression levels of *LIF* in other cerebral organoid models **C-D** GFAP staining in control and LIF-treated organoids fixed, sectioned and stained on day 90 from multiple hPSC lines showing the LIF protocol is robust and reproducible across multiple cell lines. Scale bars: 100 μm.

**Figure S3.**
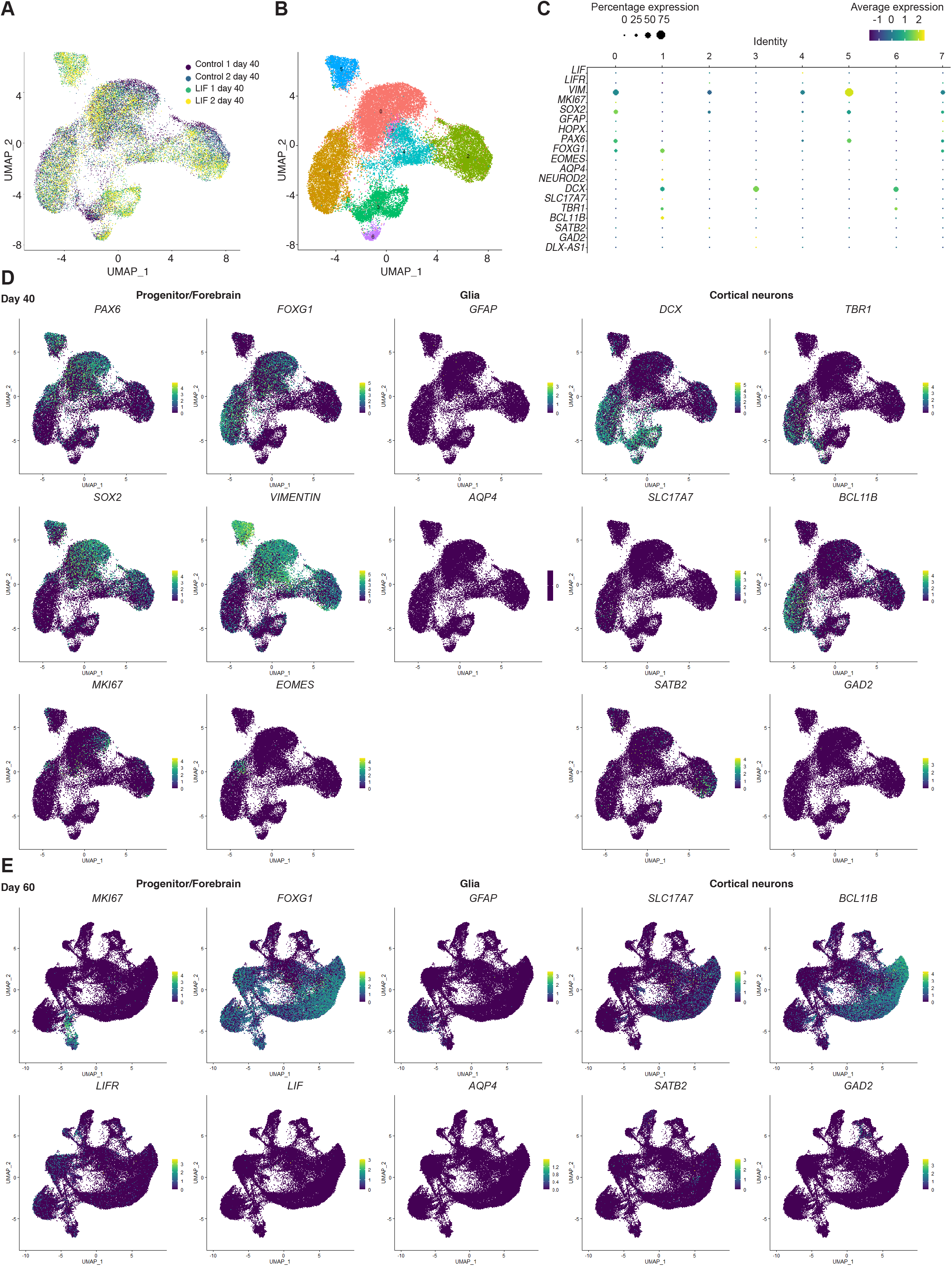
**A** scRNA-seq experiments showing plots for original identity and **B** Seurat clusters in control and LIF-treated organoids on day 40. Two independent batches of 10 organoids each were processed and analyzed by scRNA-seq for both control and LIF condition. **C** Genes selected from B to mark selected populations of interest (dividing cells, neuronal cell types, neuronal progenitors, astrocytes, postmitotic neurons and radial glial cells) in control and LIF-treated organoid clusters from day 40. **D** Feature plots depicting distribution of the expression of key selected progenitor/forebrain, glia, and cortical neuron genes in control and LIF-treated organoids on day 40. **E** Feature plots depicting the distribution of the expression of key selected progenitor/forebrain, glia, and cortical neuron genes in control and LIF-treated organoids on day 60.

**Figure S4.**
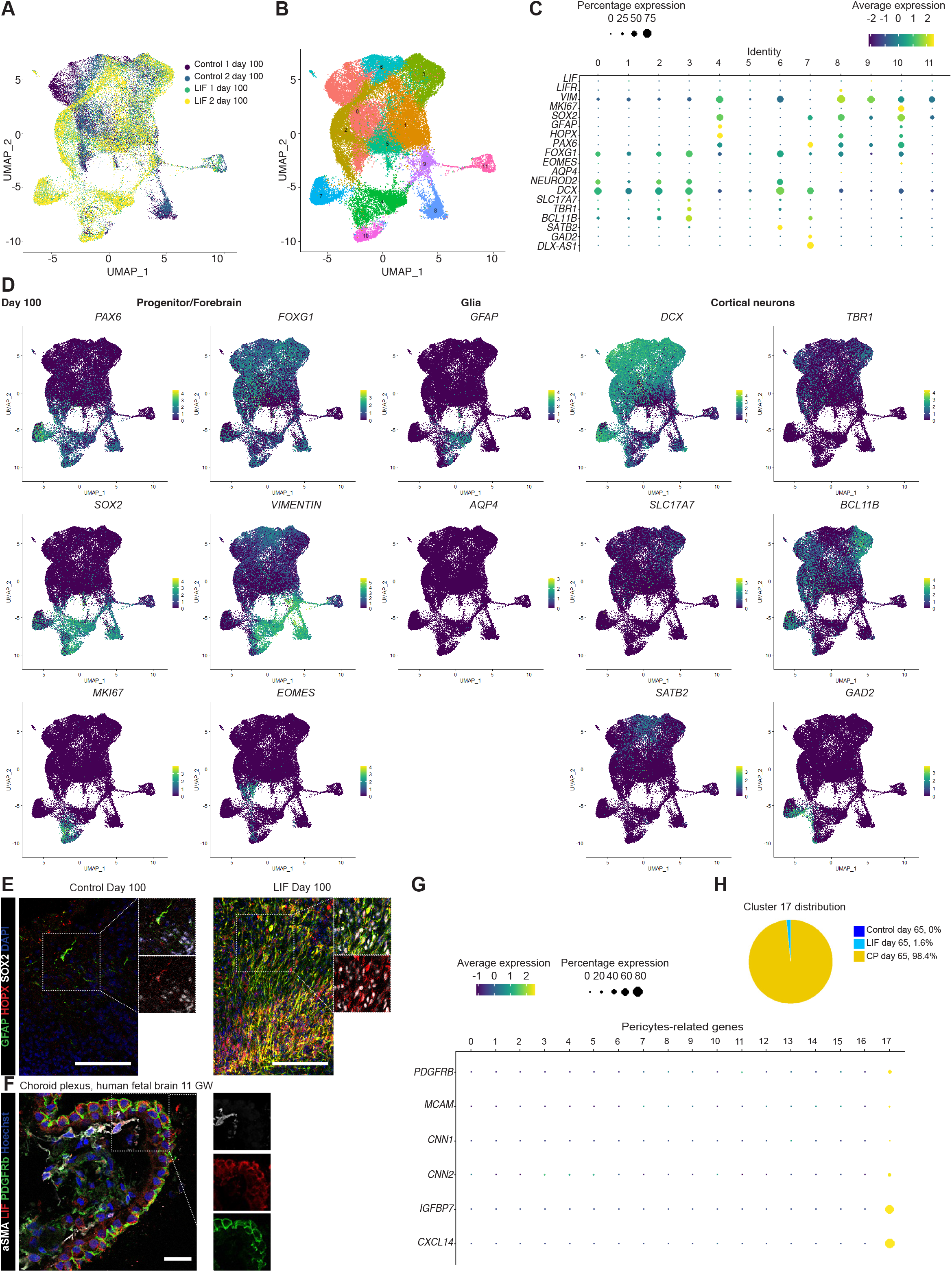
**A** scRNA-seq experiments showing plots for original identity and **B** Seurat clusters in control and LIF-treated organoids on day 100. Two independent batches of 10 organoids each were processed and analyzed by scRNA-seq for both control and LIF condition. **C** Genes selected from B to mark selected populations of interest (dividing cells, neuronal cell types, neuronal progenitors, astrocytes, postmitotic neurons and radial glial cells) in control and LIF-treated organoid clusters from day 100. **D** Feature plots depicting distribution of the expression of key selected progenitor/forebrain, glia, and cortical neuron genes in control and LIF-treated organoids on day 100. **E** GFAP staining in control (left) and LIF-treated (right) organoids that were fixed, sectioned and stained on day 100. Nuclei are stained in blue with Dapi. Scale bars:100 μm. **F** LIF, PDGFRb and aSMA staining in human fetal cortex and choroid plexus from human fetal brain 11GW sections. Nuclei are stained in blue with Dapi. Scale bars: 25 μm. **G** Genes selected from Figure 4N to mark selected key pericyte markers in cluster 17 in CP assembloids from day 65. **H** Pie chart showing cluster 17 distribution (percentages) in control organoids, LIF-treated organoids and CP assembloids on day 65.

